## Supplemental Figures and Tables for "Cryo-EM structures reveal high-resolution mechanism of a DNA polymerase sliding clamp loader"

### Supplemental Figures, Videos and Tables

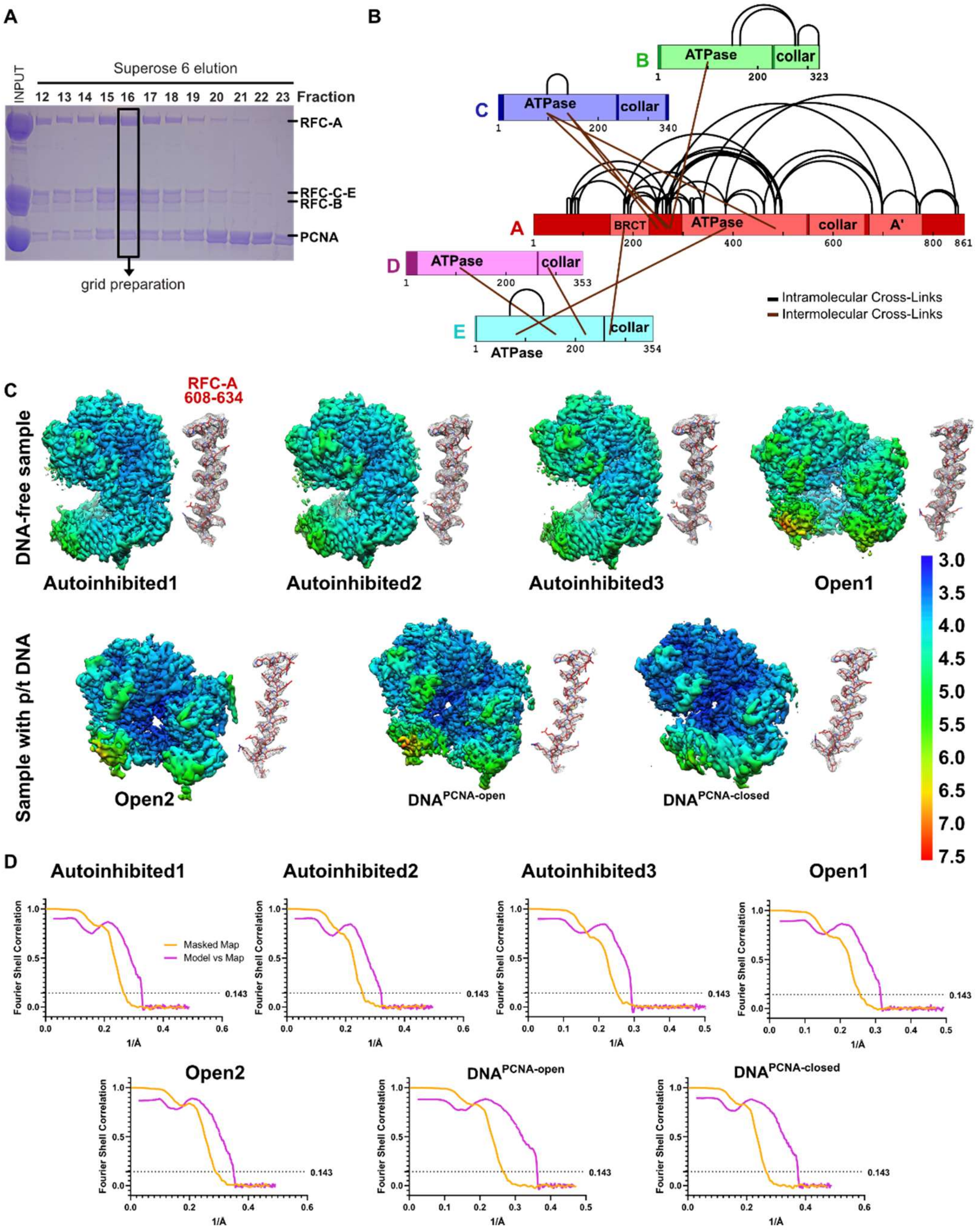


**Figure S1. Characterization and cryo-EM of full-length RFC:PCNA . (A)** SDS-PAGE gel of purified RFC and PCNA after gel filtration. A fraction with stochiometric amounts of RFC and PCNA was used for grid preparation. **(B)** Crosslinking of the RFC:PCNA at a concentration of 1 mM Bissulfosuccinimidyl suberate (BS3) led to the identification of intermolecular and intramolecular crosslinks in RFC, and are shown in schematic representation. 88% of the crosslinks mapped to RFC-A and no crosslinks in PCNA were detected, although PCNA was detected in the sample. **(C)** Local resolution of reconstructions (center) and a representative section of each complex subunit for each reconstruction. **(D)** Fourier shell correlation (FSC) curves for the two halves of the reconstructions as well as model vs map curves.

### Supplemental Figures, Videos and Tables

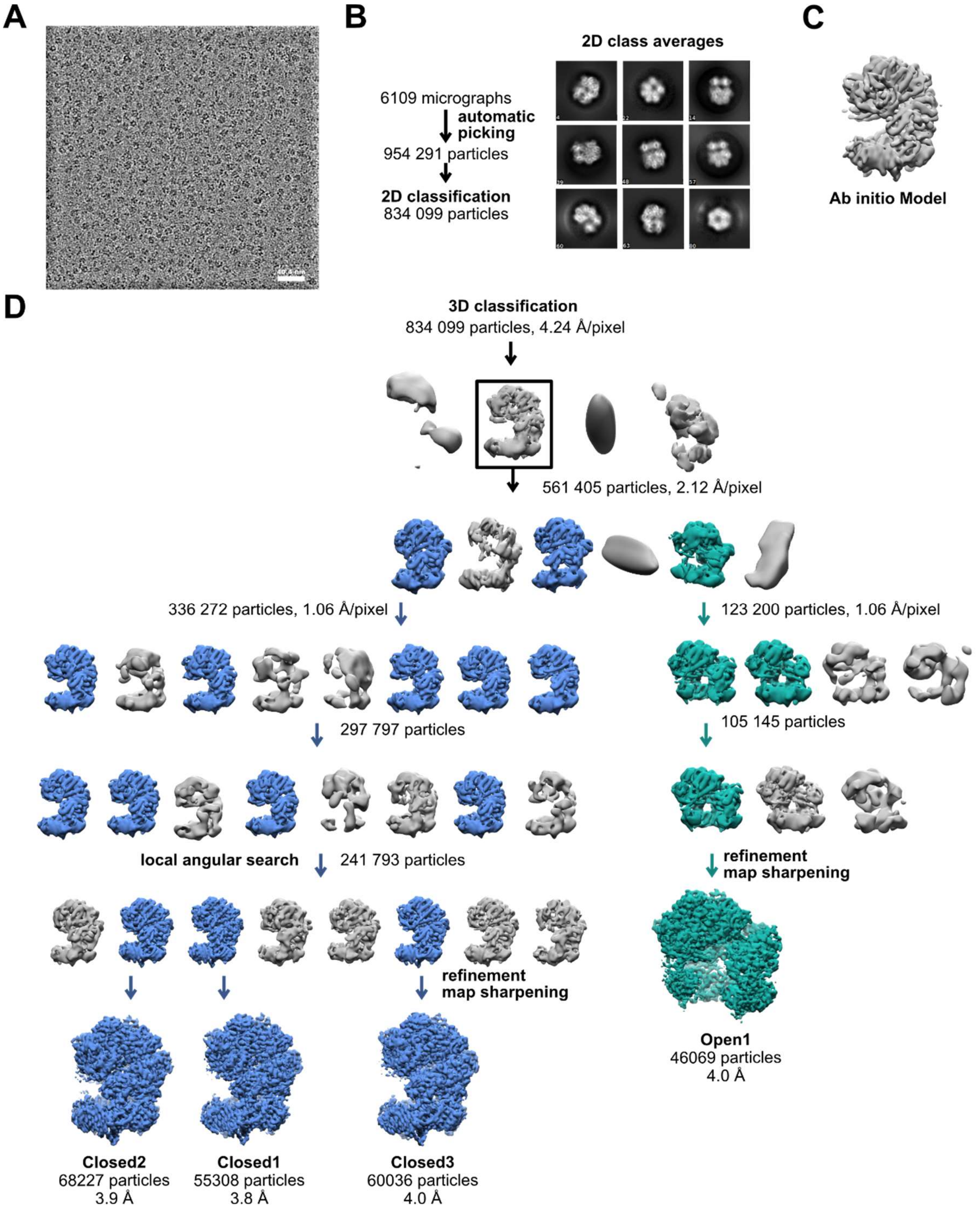


**Figure S2. Schematic of yRFC:PCNA cryo-EM processing. (A)** A down-filtered micrograph taken on a Thermo Fisher Scientific Titan Krios with a Gatan K2 detector is displayed. **(B)** 2D class averages show different side views. **(C)** The 3D reference for refinement was generated *ab initio* with cisTEM (Grant et al., 2018). **(D)** The ab initio model was downfiltered to 50 Å and used as reference for 3D classification. The first round of classification was performed with the 2x binned particle stack. Further rounds of classification with the unbinned stack improved the resolution. 3D classification with local angular search further helped to improve the resolution of the reconstructions representing complexes with closed PCNA (blue).

### Supplemental Figures, Videos and Tables

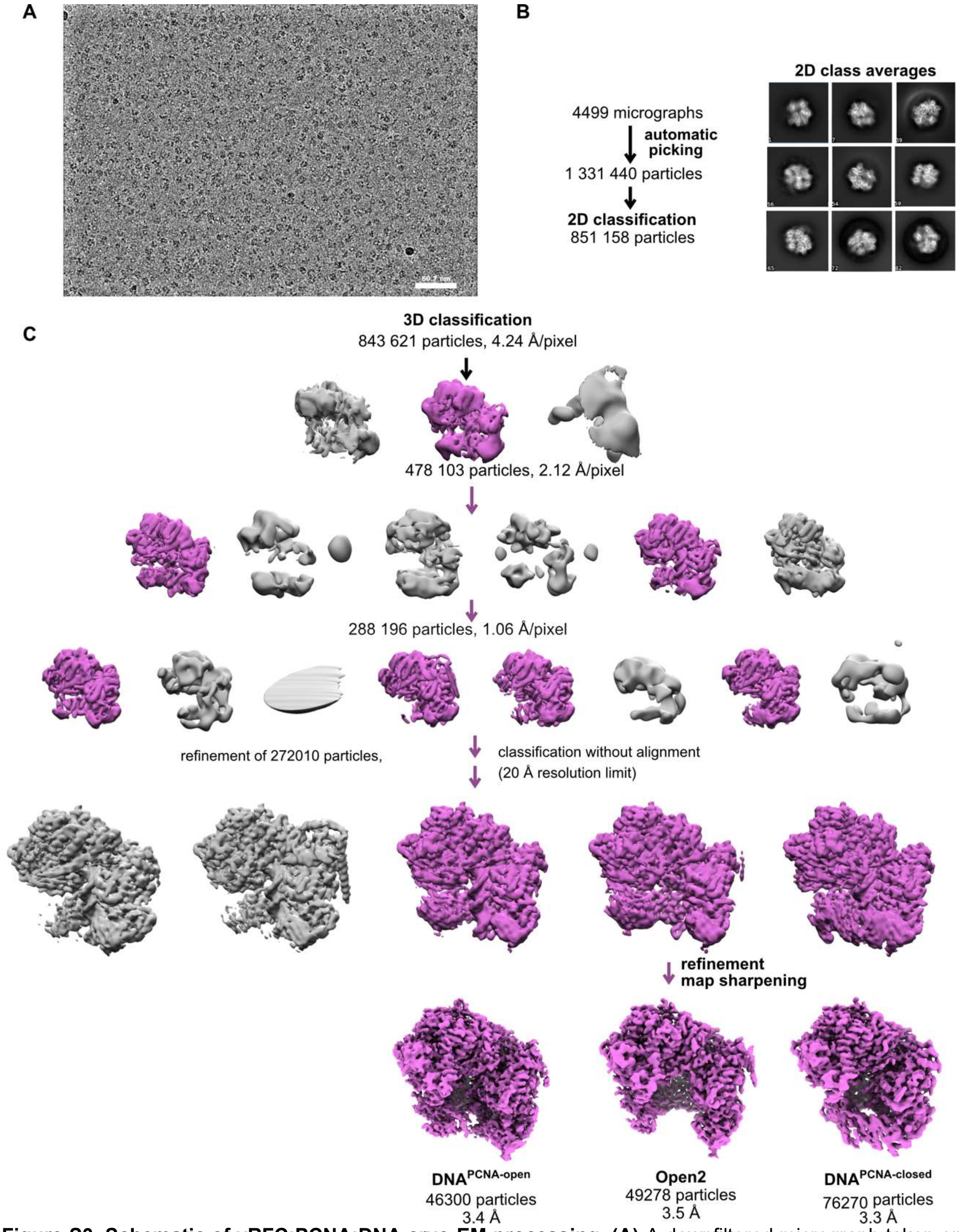


**Figure S3. Schematic of yRFC:PCNA:DNA cryo-EM processing. (A)** A downfiltered micrograph taken on a Thermo Fisher Scientific Titan Krios with a Gatan K3 detector is displayed. **(B)** 2D class averages show different side views. **(C)** Class Open1 of the dataset without DNA was downfiltered to 60 Å and used as reference for 3D classification. The first round of classification was performed with the 4x binned particle stack and the second round of classification with the 2x binned stack. Further classification with the unbinned stack with resolution limit and without alignment helped to separate different conformational states and improved the resolution.

### Supplemental Figures, Videos and Tables

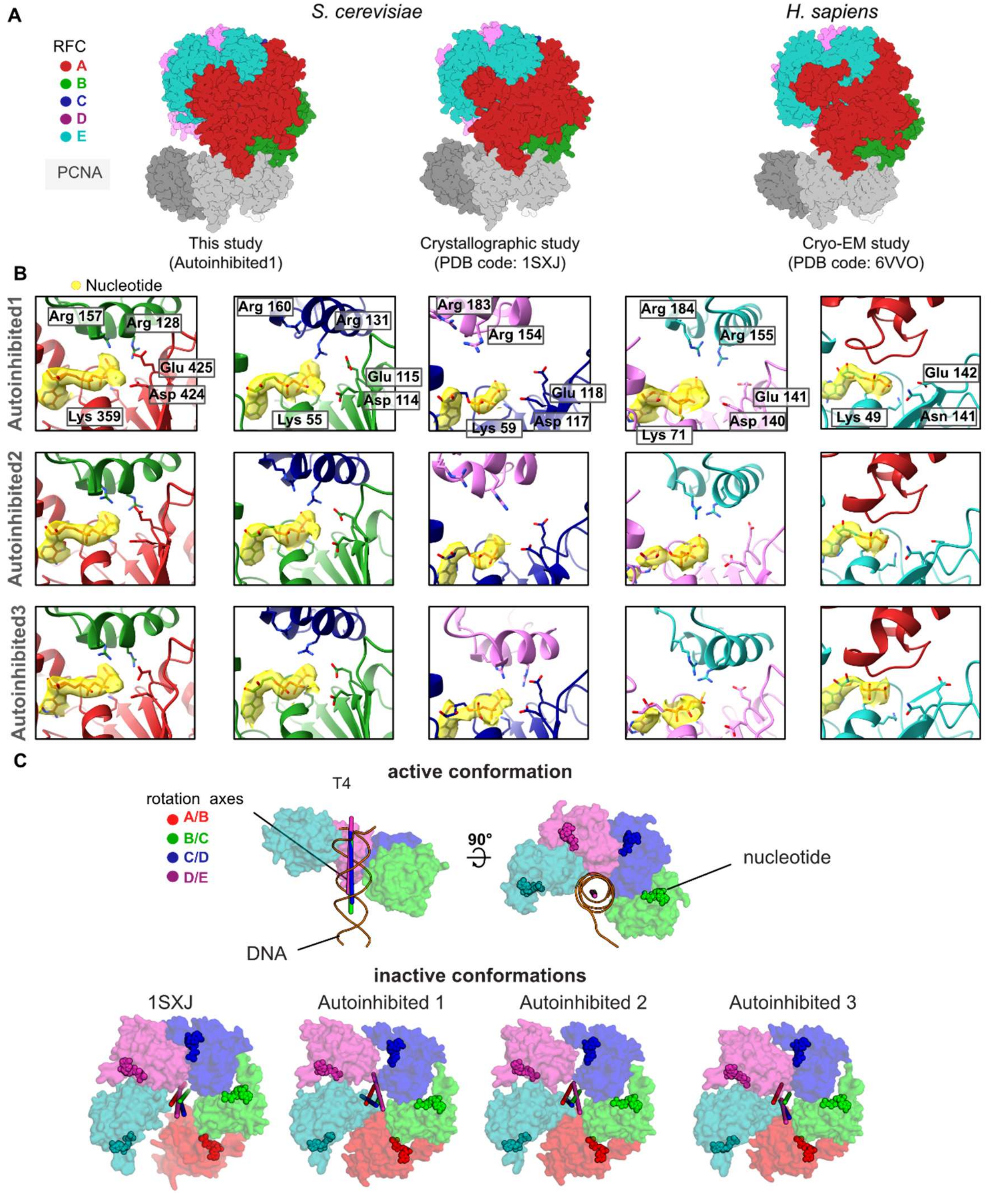


**Figure S4. RFC:PCNA complexes in autoinhibited conformations. (A)** Side view of the atomic models of Autoinhibited1, the RFC:PCNA crystal structure, and of the human RFC:PCNA complex show similarity. **(B)** Close-up of the nucleotide binding sites in Autoinhibited1-3. The cryo-EM map is shown in yellow overlaying the atomic model. The catalytically important Walker A lysine, Walker B glutamates, and *trans*-acting arginine fingers are shown. The arginine fingers are distant in RFC-B,C,D in Autoinhibited1&2, and in RFC-B,D of Autoinhibited3, rendering these active sites inactive. RFC-E is not catalytically competent and has ADP-bound. The A' domain does not donate *trans*-acting arginine fingers. **(C)** Top views on the AAA+ spiral of the T4 clamp loader, RFC:PCNA crystal structure (PDB 1SXJ) and Autoinihibited1-3. The T4 clamp loader, which has DNA bound, is in an active conformation. Here, the rotation axes that relate the subunits are coincident with each other and the central axis of DNA. In contrast, the symmetry of the AAA+ spiral of RFC in the autoinhibited conformation is distorted, and the axes are skewed in all these structures.

### Supplemental Figures, Videos and Tables

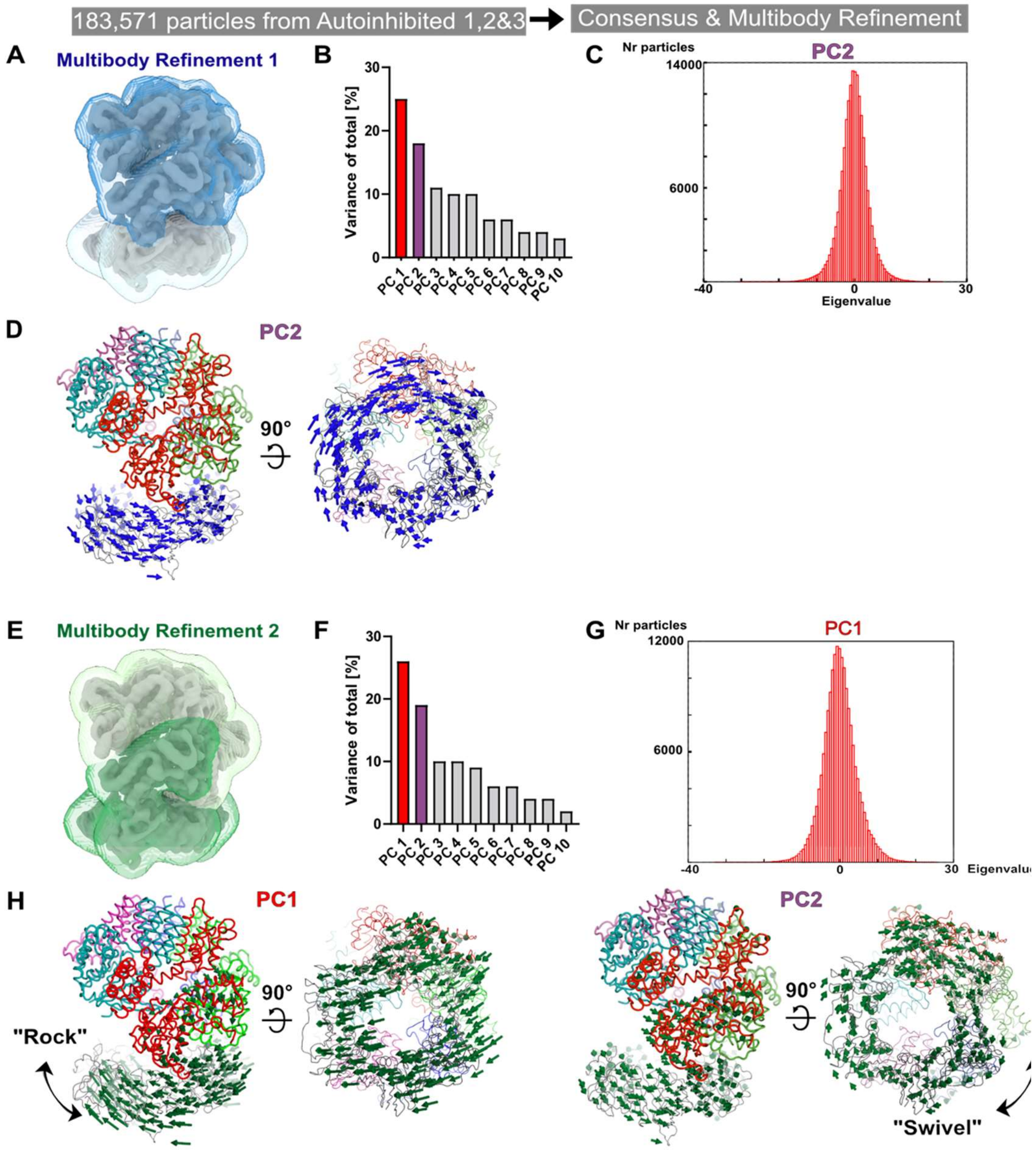


**Figure S5. Multibody analyses to investigate the dynamic initial complex of RFC with PCNA. (A)** The two masks used for the first multibody refinement define PCNA and RFC as separate rigid bodies. **(B)** Principal Component (PC) analysis revealed the two most dominant motions with RFC and PCNA. **(C)** Amplitude histogram of PC2 is unimodal**. (D)** PC2 reveals a swiveling motion. The $C_\alpha$ displacement is indicated by modevector generated arrows, scaled down by a factor of 2. **(E)** The two masks used for the second multibody refinement were chosen to match domain boundaries determined with the ENM DynOmics server (Li et al., 2017) and to capture motion between the A' and the AAA+ module of the A-gate. **(F)** PC analysis revealed two dominant motions. **(G)** Amplitude histogram of PC1 is unimodal**. (H)** Multibody refinement 2 also revealed rocking and swiveling as dominant motions.

### Supplemental Figures, Videos and Tables

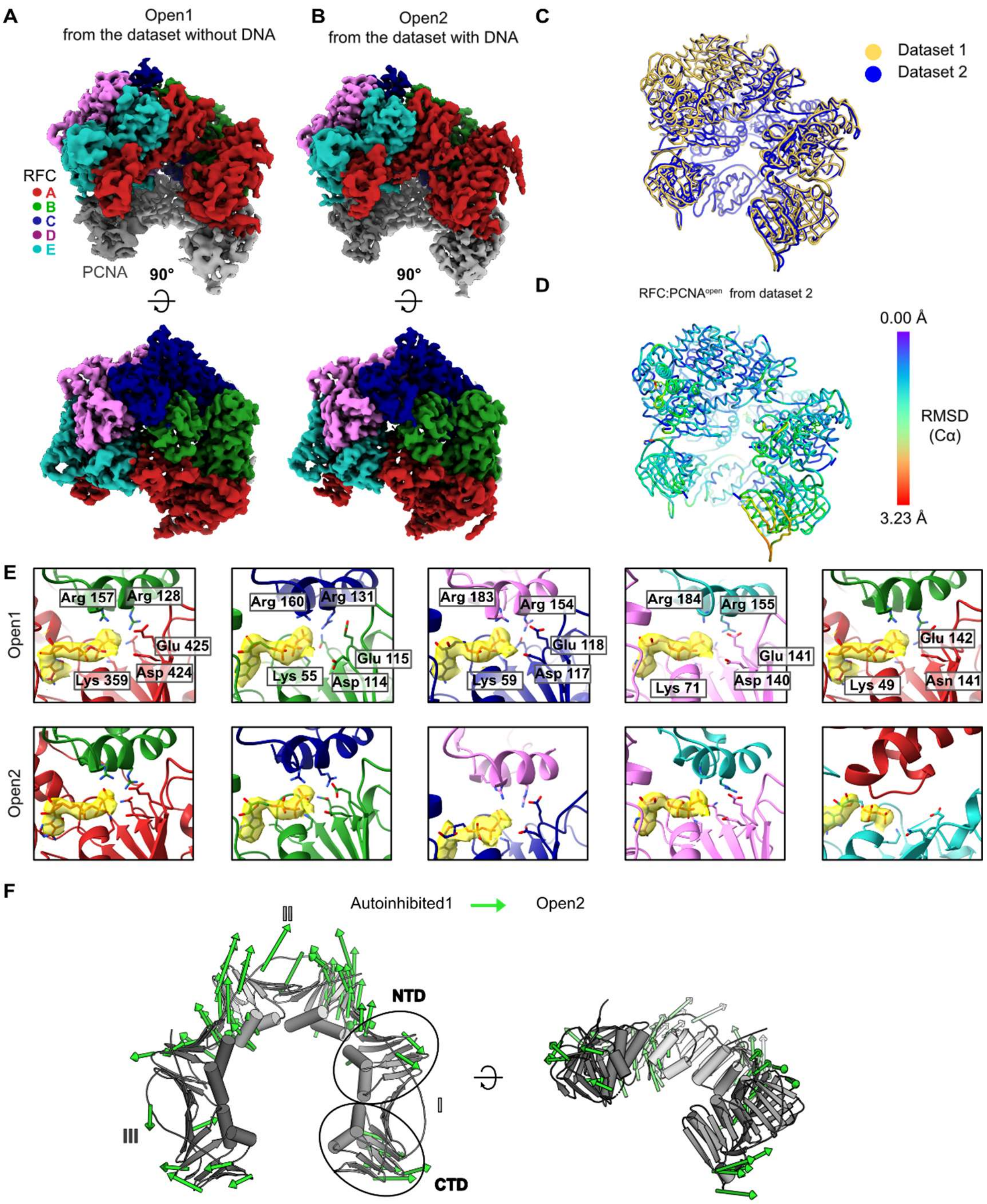


**Figure S6. Comparison of RFC bound to open PCNA from different datasets. (A)** Top and side view of the cryo-EM map of RFC bound to open PCNA, which was obtained from the dataset without DNA. **(B)** Top and side view of the cryo-EM map of RFC bound to open PCNA obtained from the dataset with DNA. **(C)** Overlay of the two models for Open1 and Open2 shows that the two models strongly resemble each other. **(D)** Open1 superposed to Open2. Open2 is colored by RMSD. **(E)** Close-up of the nucleotide binding sites in Open1 and in Open2. The cryo-EM map is shown in yellow overlaying the atomic model. Critical catalytic residues are shown as sticks. All active sites are occupied with ATPγS. (**F**) PCNA intrasubunit distortions that occur for opening. The Cα displacement is indicated by modevector-generated arrows, scaled up by a factor of four.

### Supplemental Figures, Videos and Tables

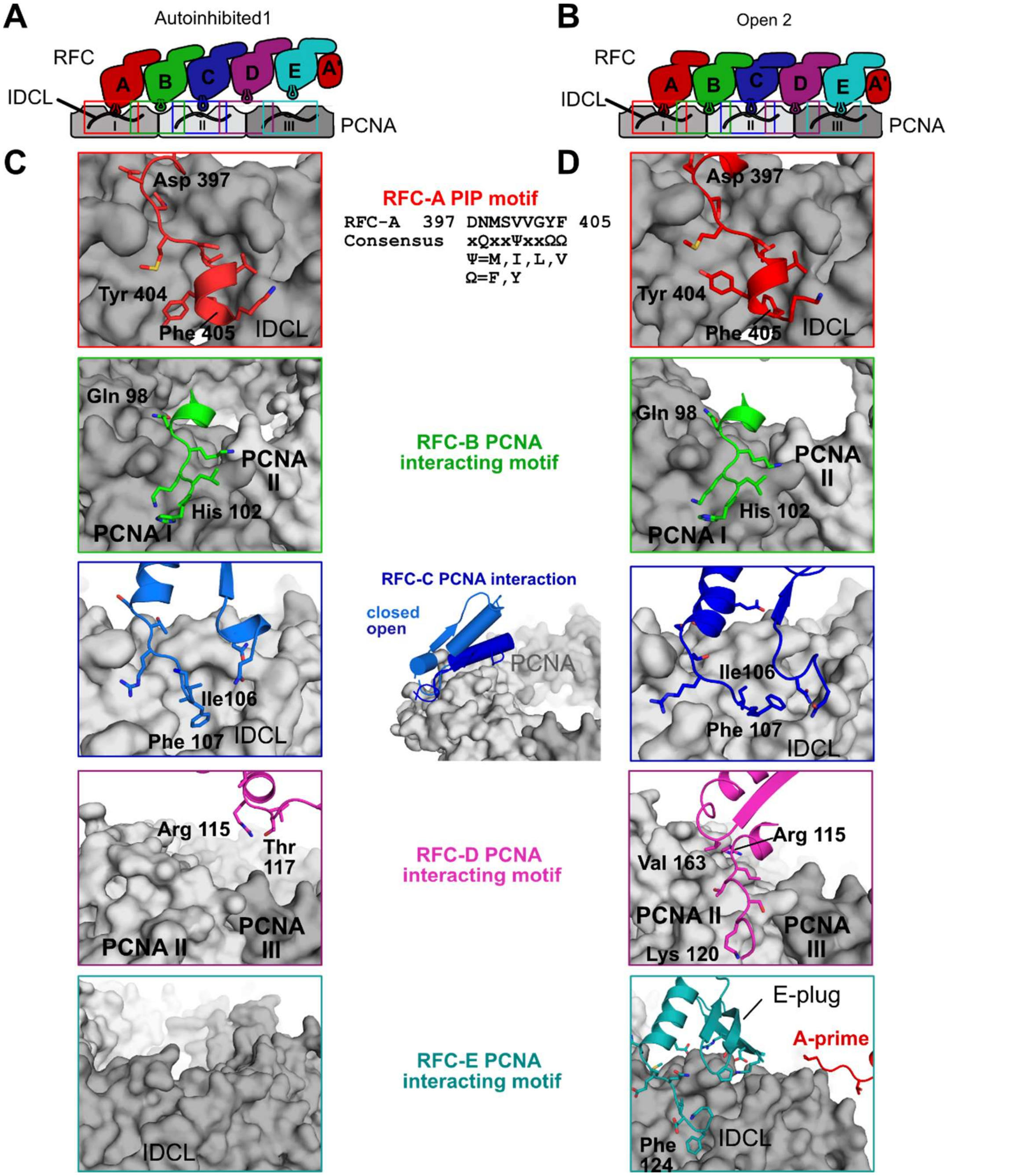


**Figure S7. RFC interacts with all five subunits to hold PCNA open. (A)** Overview of interaction sites of RFC with PCNA in the autoinhibited conformation. The three PCNA subunits and RFC's AAA+ module are shown in a cartoon flattened onto the page. RFC-D and RFC-E are not in contact with PCNA. **(B)** Cartoon overview of interaction sites of RFC with PCNA in the open conformation. RFC-D and RFC-E now contact PCNA. **(C)** Close up views of the RFC-PCNA interaction sites in Autoinhibited1 are shown, the rest is omitted for clarity. The contact between PCNA and RFC-A is mediated by a short helix and adjacent hydrophobic and aromatic residues that insert into PCNA's hydrophobic pockets. This conformation is commonly seen in binding partners which contain a PCNA-interacting protein (PIP) motif or derived motifs. **(D)** Contacts of RFC-A and RFC-B with PCNA do not change significantly upon PCNA opening. The interaction of PCNA with RFC-C becomes more extensive, and RFC-D and RFC-E establish new contacts to PCNA. RFC-C and RFC-E insert into PCNA's hydrophobic pocket but do not employ a PIP motif. The E-plug and A' domain reinforce the interaction with PCNA.

### Supplemental Figures, Videos and Tables

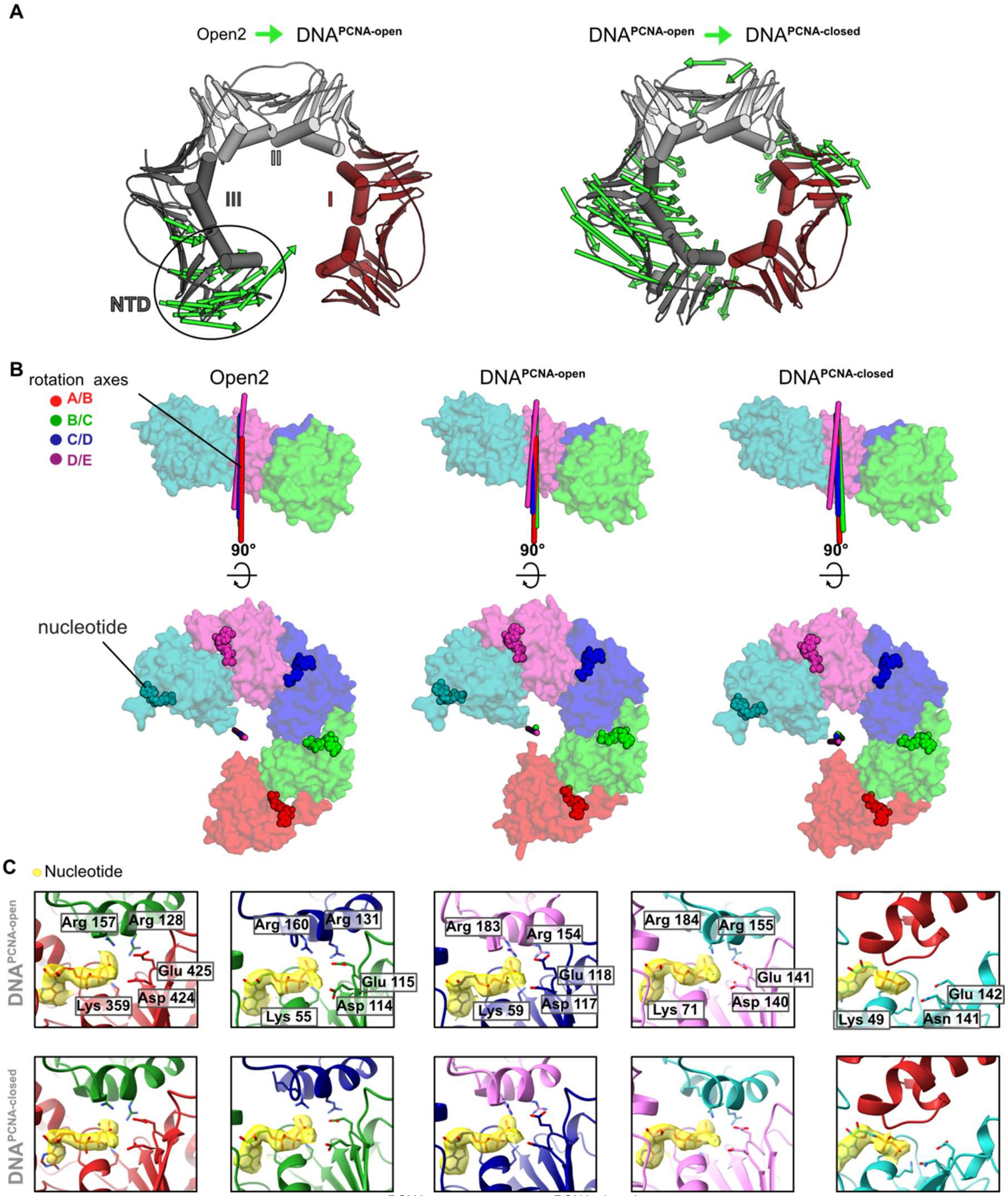


**Figure S8: (A)** PCNA constriction in DNA$^{PCNA\text{-}open}$ and DNA$^{PCNA\text{-}closed}$. Displacement vectors between Open2 and DNA$^{PCNA\text{-}open}$ are shown as green arrows, scaled by a factor of four (left). Displacement vectors between DNA$^{PCNA\text{-}open}$ and DNA$^{PCNA\text{-}closed}$ are shown as green arrows, scaled by a factor of four (right). Upon DNA binding, the PCNA lock-washer constricts in DNA$^{PCNA\text{-}open}$, due to a motion at the NTD of PCNA-III. PCNA is closed in a puckered conformation in DNA$^{PCNA\text{-}closed}$ through constricting motions of PCNA-I and PCNA-III. **(B)** The overlay of the rotation axes in the open conformation of RFC is indicative for spiral symmetry. In the DNA-bound structures (DNA$^{PCNA\text{-}open}$ and DNA$^{PCNA\text{-}closed}$), the rotation axes become more tilted, indicating that DNA binding and PCNA closure slightly disrupt the symmetric arrangement of the AAA+ spiral. **(C)** Close-up of the nucleotide binding sites in DNA$^{PCNA\text{-}open}$ and DNA$^{PCNA\text{-}closed}$. The cryo-EM map is shown in yellow overlaying the atomic model. Critical catalytic residues are shown as sticks. All active sites are occupied with ATPγS.

### Supplemental Figures, Videos and Tables

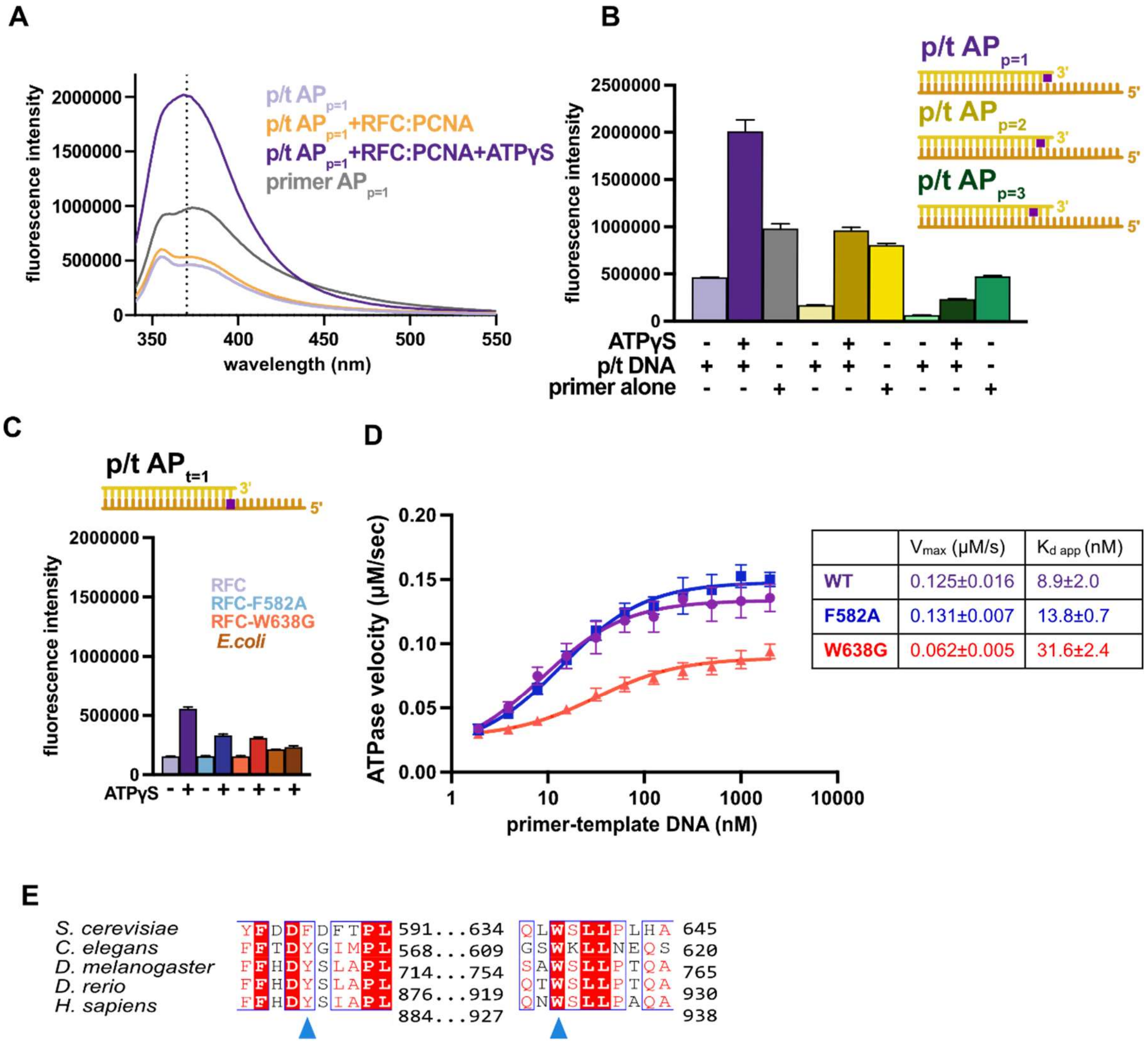


**Figure S9: The separation wedge has two critical residues. (A)** Fluorescence intensity traces for 375 nM of p/t $AP_{p=1}$ in the presence of RFC:PCNA with and without nucleotide or primer $AP_{p=1}$. **(B)** Placement of 2AP at different positions. The fluorescence with p/t $AP_{p=1}$ in the presence of ATPγS and RFC:PCNA increases ~4-fold in relation to the sample without ATPγS, whereas placement of the oligo further away from the 3'-OH end reduces the fluorescence increase to ~2-fold. **(C)** Results from (Figure 6 B) could be recapitulated using p/t-DNA with 2-AP in the template strand (t=1). The *E. coli* clamp loader, which does not have a separation pin, does not change fluorescence in the presence of ATPγS. **(D)** ATPase activity of the 'separation pin' mutants. DNA binding affinity and maximum ATP hydrolysis rate is reduced in $RFC^{W638G}$. **(E)** The 'separation pin' is conserved among eukaryotes. Sequence alignment shows the conservation of the 'separation pin' among 5 eukaryotic species. The conserved sequences are marked by blue boxes. The fully conserved residues are in white with a red background, the highly conserved residues are in red, and the less conserved ones are in black. F582 and W638 are pointed out by the blue arrow.

### Supplemental Figures, Videos and Tables

**A**

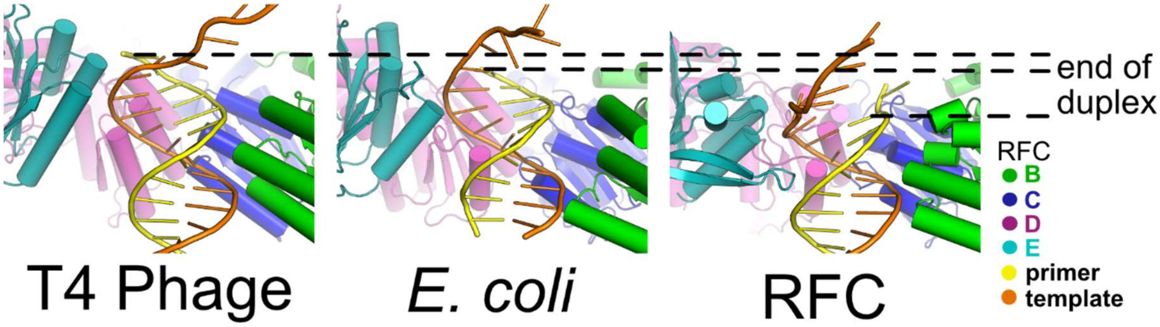


**B**

| Clamploader interactions with p/t-DNA | | | |
|---|---|---|---|
| | Interaction Area B, C, D, E subunits (Å$^2$) | Interaction Area A subunit (Å$^2$) | Reference |
| T4 Phage | 1149 | 659 | Kelch et al 2011 |
| *E. coli* | 1100 | 619 | Simonetta et al 2009 |
| *S. cerevisiae* | 762 | 1369 | this work |

**Figure S10: Differences in how duplex p/t-DNA is held in the central chamber of clamp loaders. (A)** In the T4 and *E.* coli clamp loaders, the duplex p/t-DNA extends farther into the central chamber, enabling more substantial contacts with the B-E subunits compared to RFC. **(B)** Contribution of RFC subunits to DNA binding. RFC-A dominates contact with p/t-DNA when compared to other clamp loaders.

**VIDEOS**

**Video S1** RFC Motion along Eigenvalue 1 with masks on PCNA and RFC
**Video S2** RFC Motion along Eigenvalue 2 with masks on PCNA and RFC
**Video S3** Morph closed to open
**Video S4** PCNA loading by RFC

### Supplemental Figures, Videos and Tables

#### Table S1 List of BS3 Crosslinks

| XlinkX Score | Type | # Crosslink Spectral Matches | Sequence A | Position A | Sequence B | Position B | Protein A | Protein B |
|---|---|---|---|---|---|---|---|---|
| 58.66 | Inter | 1 | [K]LHLPPGK | 100 | [K]LAATR | 274 | RFC4 | RFC1 |
| 58.64 | Inter | 3 | [K]LELNVVSSPYHLEITPSDMGNNDR | 82 | S[K]TLLNAGVK | 385 | RFC5 | RFC1 |
| 56.99 | Inter | 3 | [K]YVNTFMK | 285 | DIL[K]R | 220 | RFC2 | RFC5 |
| 56.47 | Inter | 1 | NQI[K]DFASTR | 98 | [K]LAATR | 274 | RFC3 | RFC1 |
| 52.59 | Inter | 2 | E[K]VKNFAR | 109 | TME[K]YSK | 160 | RFC2 | RFC5 |
| 50.97 | Inter | 1 | NQI[K]DFASTR | 98 | RPDANSI[K]SR | 484 | RFC3 | RFC1 |
| 48.17 | Inter | 1 | GASEALA[K]R | 182 | [K]IVKER | 269 | RFC1 | RFC5 |
| 45.16 | Inter | 1 | YT[K]NTR | 139 | [K]EEER | 267 | RFC3 | RFC1 |
| 41.65 | Inter | 1 | [K]LEEQHNIATK | 249 | YT[K]NTR | 139 | RFC1 | RFC3 |
| 91.6 | Intra | 3 | [K]LEEQHNIATK | 249 | RPDANSI[K]SR | 484 | RFC1 | RFC1 |
| 72.73 | Intra | 4 | EAELLV[K]KEEER | 266 | [K]LAATR | 274 | RFC1 | RFC1 |
| 71.87 | Intra | 12 | QLIAGMPAEGGDGEAAE[K]AR | 245 | R[K]LEEQHNIATK | 249 | RFC1 | RFC1 |
| 71.27 | Intra | 2 | E[K]FKLDPNVIDR | 495 | [K]LAATR | 274 | RFC1 | RFC1 |
| 71.03 | Intra | 1 | F[K]LDPNVIDR | 497 | [K]LAATR | 274 | RFC1 | RFC1 |
| 71.03 | Intra | 9 | [K]TSTPLILICNER | 446 | S[K]TLLNAGVK | 385 | RFC1 | RFC1 |
| 64 | Intra | 1 | EAELLV[K]KEEER | 266 | S[K]KLAATR | 273 | RFC1 | RFC1 |
| 62.71 | Intra | 1 | RPDANSI[K]SR | 484 | SA[K]YYR | 678 | RFC1 | RFC1 |
| 62.2 | Intra | 2 | YAPTNLQQVCGN[K]GSVMK | 314 | L[K]NWLANWENSKK | 321 | RFC1 | RFC1 |
| 61.3 | Intra | 4 | EAELLVK[K]EEERSK | 267 | [K]LAATR | 274 | RFC1 | RFC1 |
| 60.15 | Intra | 1 | FAFACNQSN[K]IIEPLQSR | 149 | VT[K]NLAQVK | 275 | RFC4 | RFC4 |
| 60.15 | Intra | 3 | YS[K]LSDEDVLKR | 165 | VT[K]NLAQVK | 275 | RFC4 | RFC4 |
| 58.98 | Intra | 1 | IPATV[K]SGFTR | 767 | HAG[K]DGSGVFR | 340 | RFC1 | RFC1 |
| 58.55 | Intra | 4 | GASEALA[K]R | 182 | VT[K]SISSK | 190 | RFC1 | RFC1 |
| 57.1 | Intra | 3 | RPDANSI[K]SR | 484 | [K]EEER | 267 | RFC1 | RFC1 |
| 56.99 | Intra | 1 | KLEEQHNIAT[K]EAELLVK | 259 | [K]EEER | 267 | RFC1 | RFC1 |
| 56.99 | Intra | 1 | DNVVREED[K]LWTVK | 296 | [K]EEER | 267 | RFC1 | RFC1 |
| 56.41 | Intra | 1 | [K]YNSMTHPVAIYR | 773 | LGTSTD[K]IGLR | 698 | RFC1 | RFC1 |
| 53.33 | Intra | 1 | Y[K]CVIINEANSLTK | 136 | L[K]IDVR | 69 | RFC5 | RFC5 |
| 52.59 | Intra | 2 | [K]ASSPTVKPASSK | 77 | [K]TKPSSK | 90 | RFC1 | RFC1 |
| 52.59 | Intra | 2 | HAG[K]DGSGVFR | 340 | GSVM[K]LK | 319 | RFC1 | RFC1 |
| 52.59 | Intra | 2 | ASSPTV[K]PASSK | 84 | [K]TKPSSK | 90 | RFC1 | RFC1 |
| 51.79 | Intra | 2 | [K]LEEQHNIATK | 249 | [K]LAATR | 274 | RFC1 | RFC1 |
| 50.97 | Intra | 1 | [K]TATSKPGGSK | 845 | S[K]TLLNAGVK | 385 | RFC1 | RFC1 |
| 50.34 | Intra | 1 | KMPVSNVIDVSETPEGE[K]K | 68 | LPLPA[K]R | 75 | RFC1 | RFC1 |
| 49.59 | Intra | 4 | EKF[K]LDPNVIDR | 497 | RPDANSI[K]SR | 484 | RFC1 | RFC1 |
| 47.92 | Intra | 1 | LGTSTD[K]IGLR | 698 | [K]LAATR | 274 | RFC1 | RFC1 |
| 47.92 | Intra | 1 | S[K]TLLNAGVK | 385 | [K]LAATR | 274 | RFC1 | RFC1 |
| 47.92 | Intra | 1 | GASEALA[K]R | 182 | [K]LAATR | 274 | RFC1 | RFC1 |
| 47.85 | Intra | 2 | SISS[K]TSVVVLGDEAGPK | 195 | [K]LEEQHNIATK | 249 | RFC1 | RFC1 |
| 47.85 | Intra | 1 | [K]YNSMTHPVAIYR | 773 | [K]TATSKPGGSK | 845 | RFC1 | RFC1 |
| 46.57 | Intra | 4 | R[K]LEEQHNIATK | 249 | GASEALA[K]R | 182 | RFC1 | RFC1 |
| 46.35 | Intra | 1 | [K]ASSPTVKPASSK | 77 | VT[K]SISSK | 190 | RFC1 | RFC1 |
| 45.16 | Intra | 1 | YAPTNLQQVCGN[K]GSVMK | 314 | [K]EEER | 267 | RFC1 | RFC1 |
| 45.16 | Intra | 1 | E[K]FKLDPNVIDR | 495 | [K]EEER | 267 | RFC1 | RFC1 |
| 45.16 | Intra | 2 | [K]LEEQHNIATK | 249 | [K]EEER | 267 | RFC1 | RFC1 |
| 44.72 | Intra | 1 | E[K]FKLDPNVIDR | 495 | RPDANSI[K]SR | 484 | RFC1 | RFC1 |
| 44.45 | Intra | 1 | YAPTNLQQVCGN[K]GSVMK | 314 | [K]LEEQHNIATK | 249 | RFC1 | RFC1 |
| 44.14 | Intra | 2 | NLP[K]MRPFDR | 462 | S[K]TLLNAGVK | 385 | RFC1 | RFC1 |
| 44.14 | Intra | 1 | RPDANSI[K]SR | 484 | GASEALA[K]R | 182 | RFC1 | RFC1 |
| 44.12 | Intra | 1 | [K]LEEQHNIATK | 249 | [K]TKPSSK | 90 | RFC1 | RFC1 |
| 43.7 | Intra | 1 | NLP[K]MRPFDR | 462 | LGTSTD[K]IGLR | 698 | RFC1 | RFC1 |
| 43.7 | Intra | 1 | [K]YNSMTHPVAIYR | 773 | TATS[K]PGGSK | 850 | RFC1 | RFC1 |
| 41.98 | Intra | 2 | LGTSTD[K]IGLR | 698 | RPDANSI[K]SR | 484 | RFC1 | RFC1 |
| 41.98 | Intra | 1 | [K]LEEQHNIATK | 249 | F[K]LDPNVIDR | 497 | RFC1 | RFC1 |
| 41.98 | Intra | 1 | HAG[K]DGSGVFR | 340 | VT[K]SISSK | 190 | RFC1 | RFC1 |
| 41.94 | Intra | 1 | NQI[K]DFASTR | 98 | YT[K]NTR | 139 | RFC3 | RFC3 |
| 40.95 | Intra | 1 | NLAQV[K]ESVR | 281 | IHKLNN[K]A | 322 | RFC4 | RFC4 |
| 40.92 | Intra | 1 | KLPLPA[K]R | 75 | [K]EEER | 267 | RFC1 | RFC1 |

**Supplemental Figures, Videos and Tables**

**Table S2 Cryo-EM data collection, processing, and model statistics**

| Dataset | **NO DNA** | | | | | **DNA** | |
|---|---|---|---|---|---|---|---|
| Magnification | 130,000 | | | | | 81,000 | |
| Voltage (keV) | 300 | | | | | 300 | |
| Cumulative exposure (e–/Å 2 ) | 49-51 | | | | | 40 | |
| Detector | K2 Summit | | | | | K3™ | |
| Pixel size (Å) | 1.059 | | | | | 1.06 | |
| Defocus range (µm) | -1.1 to -2.4 | | | | | -1.2 to -2.3 | |
| Micrographs used (no.) | 6109 | | | | | 4499 | |
| Initial particle images (no.) | 954,291 | | | | | 1,331,440 | |
| Symmetry | C1 | | | | | | |
| **Class Name** | **yRFC:PCNA Autoinhibited1** | **yRFC:PCNA Autoinhibited2** | **yRFC:PCNA Autoinhibited3** | **yRFC:PCNA Open1** | **yRFC:PCNA Open2** | **yRFC:PCNA DNA-open** | **yRFC:PCNA DNA-closed** |
| Final Refined particles (no.) | 55,308 | 68,227 | 60,036 | 46,069 | 63,752 | 46300 | 76270 |
| Applied B factor ($Å^2$) | -100 | -159.352 | -163.938 | -100 | -106.457 | -105.857 | -105.313 |
| Map resolution (Å, FSC 0.143) | 3.8 | 3.9 | 4.0 | 4.0 | 3.5 | 3.4 | 3.3 |
| Model-Map CC_mask | 0.80 | 0.79 | 0.78 | 0.80 | 0.82 | 0.79 | 0.81 |
| Bond lengths (Å), angles (°) | 0.005, 0.668 | 0.004, 0.613 | 0.004, 0.621 | 0.005, 0.706 | 0.003, 0.562 | 0.003, 0.682 | 0.003, 0.548 |

### Supplemental Figures, Videos and Tables

| | | | | | | | |
|---|---|---|---|---|---|---|---|
| Ramachandran Outliers, Allowed, Favored | 0.00,<br>4.7, 95.3 | 0.00,<br>4.26, 95.74 | 0.00,<br>4.68, 95.32 | 0.00,<br>5.65, 94.35 | 0.00,<br>3.85, 96.15 | 0.00,<br>3.91, 96.09 | 0.00,<br>2.82, 97.18 |
| Poor rotamers (%),<br>MolProbity score, Clashscore (all atoms) | 0.00<br>1.85,<br>10 | 0.04,<br>1.85,<br>10.92 | 0.00,<br>1.89,<br>11.03 | 0.00,<br>2.01,<br>12.36 | 1.66,<br>1.94,<br>9.60 | 0.04,<br>1.83,<br>11.11 | 1.18,<br>1.71,<br>9.69 |
| Accession number,<br>EMDB, PDB | | | | | | | |

### Supplemental Figures, Videos and Tables
